# T-cell Multiomic Analysis Identifies Subsets and Mechanisms of Interaction with Epithelial Cells in Idiopathic Pulmonary Fibrosis

**DOI:** 10.1101/2025.11.19.689037

**Authors:** Ana P.M. Serezani, Julia M.R. Bazzano, Bruno D. Pascoalino, Ludmilla da Silva, Abigail J. Dietrich, Chase J. Taylor, Taylor Sherrill, Carla L. Calvi, Paula I Gonzalez-Ericsson, Erin M Wilfong, Matthew Bacchetta, Ciara M. Shaver, Lorraine B. Ware, Margaret L. Salisbury, Luc Van Kaer, Nicholas E. Banovich, Jonathan A Kropski, Timothy S Blackwell

## Abstract

Idiopathic pulmonary fibrosis (IPF) is a fatal interstitial lung disease characterized by progressive scarring and respiratory failure, with a median survival of 3–5 years. While T-cell numbers are elevated in IPF lungs, their contributions to fibrosis beyond inflammation remain poorly understood. Here, we performed multiplex imaging and single-cell RNA and protein profiling on ∼90,000 CD3⁺ T-cells from control and fibrotic lungs, revealing eleven distinct subsets of CD4⁺ and CD8⁺ T-cells. Among these, we identified a rare CD56⁺ regulatory T-cell subset that is highly activated in fibrosis and exhibits a sustained immunosuppressive phenotype. CXCR4/MIF signaling emerged as a central axis mediating T-cell–epithelial interactions, while epidermal growth factor receptor (EGFR) and TGFβ pathways dominated in multiple T-cell subsets. Our findings demonstrate that T-cells in IPF adopt nonclassical activation patterns, driven by epithelial interactions and the fibrotic microenvironment. These studies provide a foundation for exploring novel therapeutic strategies in IPF lungs by modulating T-cell behavior and communication networks.

## Introduction

Despite growing evidence that T-cells play important roles in a wide range of chronic fibrotic disorders, including systemic autoimmune rheumatic disease (SARD-ILD)^1,2,3^, their role and functions in idiopathic pulmonary fibrosis (IPF) remain unresolved. In patients with IPF, T-cell numbers are elevated in bronchoalveolar lavage fluid (BAL) and lung tissues, and both CD4 and CD8 T-cell subsets are found in greater abundance in regions of advanced fibrosis compared to areas with less severe fibrosis ^4–7^. In a previous study, we found that resident memory T-cells (T_RM_), which are commonly located within epithelial barriers of mucosal tissues ^8–10^, are significantly elevated in IPF lungs compared to controls ^11^. These findings suggest that interactions between T-cells and epithelial cells may be one of the major ways T-cells contribute to fibrosis.

The lungs typically harbor various subtypes of T-cells that are essential for maintaining the integrity of epithelial barriers ^12,13,14^. During fibrosis, alveolar epithelial cells (AECs) undergo repeated injury with incomplete repair, resulting in prolonged activation of fibroblasts with accumulation of collagen and extracellular matrix. This process, which is influenced by factors such as environmental exposures, aging, and genetic variations ^15,16,17^, also results in emergence of aberrant epithelial cell populations ^18,19^, increased apoptosis and cellular senescence ^20–23^. The role of T-cells within this environment, and specifically their influence on aberrant epithelial cells, remains poorly understood. Elucidating these interactions could offer critical insights into the underlying mechanisms of fibrosis. In this study, we sought to investigate T-cell subsets that are differentially activated in fibrotic lungs and to define their interactions with epithelial cells in the lung parenchyma.

Emerging single-cell technologies provide promising opportunities to characterize transient cellular interactions to understand the pathogenesis of diseases ^24,25^. We utilized multiplex immunofluorescence by CODEX (Co-Detection by indexing) together with Cellular Indexing of Transcriptomes and Epitopes by Sequencing (CITE-seq) of lung-isolated T-cells to evaluate the subsets with relevant activation in the lungs of IPF patients compared to controls; in parallel, we compared the activation profile of these subsets in IPF and other forms of ILD. Together, these studies provide a deeper characterization of T-cells in fibrosis and reveal how proximity and interactions with the epithelium are central to shape T-cell activation in IPF lungs.

## Methods

### Subjects and samples

Tissue samples were obtained from fibrotic lungs [Idiopathic Pulmonary Fibrosis (IPF), Interstitial Pneumonia with Autoimmune Features (IPAF), and Systemic Autoimmune Rheumatic Disease-associated Interstitial Lung Disease (SARD-ILD)] removed at the time of lung transplant surgery at Vanderbilt University Medical Center or non-fibrotic control lungs (declined donors) obtained at the time of organ procurement (**Supplemental Table 1**). The fibrotic diagnoses were determined according to the American Thoracic Society/European Respiratory Society consensus criteria as previously described ^23^. These studies were approved by the Institutional Review Board at Vanderbilt (IRB#171657, #162138, 060165).

### Multiplex immunofluorescence in lung tissue

Tissue microarray (TMA) sections were generated from paraffin-embedded lung samples of non-diseased controls (n = 6) and IPF patients (n = 6). Sections were cut at a thickness of 5 μm, placed on poly-lysine–treated coverslips, and incubated in antigen retrieval buffer (Dako Target Retrieval Solution). An additional section was stained with hematoxylin and eosin. Coverslips were incubated overnight with 14 pre-conjugated barcode antibody cocktails (Akoya Biosciences). Slides were placed on the Keyence BZ-X810 microscope stage insert, connected to the PhenoCycler automated fluidics system. Up to three fluorophore-conjugated reporters were added per cycle for hybridization with the antibodies and 4′,6-diamidino-2-phenylindole (DAPI) was used to label nuclei. Images were captured before de-hybridization, and image files were overlaid to form a large region of interest. Qupath^26^ was used to segment and annotate cell types. Cells were identified based on marker expression: pan-cytokeratin (epithelial cells), CD4 (CD4 T-cells), FOXP3 (T_REG_), and CD8 (CD8 T-cells).

### Isolation of single cell suspensions from lungs

Lung tissue biopsies were washed in PBS, minced, and frozen in RPMI media containing 10% DMSO until analysis. Minced lungs were thawed at 37°C for 3 minutes and digested in an enzymatic cocktail [Collagenase XI and DNAse I, type IV (Sigma)^27^ for CITE-seq, or collagenase I/dispase II for scRNA-seq] ^23^ and using a gentleMACS Octo Dissociator (Miltenyi Inc.) at 37°C and filtered using 100-μm sterile filters (Fisher).

### Cytometry by time of flight (CyTOF)

Peripheral blood [n=25 IPF (average age 68 ± 7.4); n=20 controls (average age 66 ± 9.3)] and lung-isolated cells [n=12 IPF and n=2 other ILD (average age 48 ± 7.6); n=16 controls (average age 33 ± 17)] were processed for CyTOF as previously described^11^. Raw data were normalized using CyTOF software and further analyzed with FlowJo version 10.10.0. Live cells were identified by excluding cisplatin-positive cells, and viSNE maps were generated using selected CD45^+^CD3^+^ live cells. Subsequently, CD4^+^CD25^+^ cells were gated, and the percentages of CD56-positive and CD56-negative cells (in peripheral blood) or CD38-positive and CD38-negative cells (in lungs) were determined for each sample.

### Cellular Indexing of Transcriptomes and Epitopes (CITE-seq)

Single-cell RNA sequencing/ CITE-seq was performed using single-cell suspensions derived from cryopreserved minced lung tissue of control and fibrotic samples. Whole lung cell suspensions (up to 1 million cells) were incubated with an Fc receptor blocking reagent (TrueStain FcX, BioLegend, USA) for 15 minutes, followed by staining with a panel of flow cytometry antibodies [anti-CD45 (APC-Cy7), anti-CD3 (PE-Cy5), and fourteen oligo-conjugated TotalSeq-C human antibodies (**Supplemental Table 2**, BioLegend)] for 30 minutes. Cells were washed three times with PBS containing 1% BSA and stained with a viability dye (Zombie Aqua kit, BioLegend). Up to 25,000 viable CD45⁺CD3⁺ cells were sorted per sample using a BD FACS ARIA and submitted for sequencing on the 10x Genomics Chromium platform. Briefly, 10,000 cells were loaded onto a 10x Chromium single-cell encapsulation chip, and libraries were prepared according to the manufacturer’s instructions. Sequencing was performed on Illumina platforms at a depth of at least 80 million reads per library.

### Analysis of sequencing data

Following alignment, demultiplexing (CellRanger-6.0.0/6.1.1), and quality-control filtering, we removed genes expressed in fewer than 3 cells, as well as cells with >L10% mitochondrial mRNA mapped UMIs, < 200 identified genes, > 5000 identified genes, total RNA counts < 1500, or total RNA counts >15000. Multiomodal data (gene expression and protein expression) were integrated using a Muon/Scanpy/Harmony/Multiomic Factor Analysis (MOFA)-based workflow which jointly utilized protein and RNA results for manual cell annotation. Whole-lung scRNA-seq from the same subjects who underwent T-cell profiling (previously reported in GSE135893 and/or GSE227136) were reanalyzed using a similar Scanpy-based workflow including label transfer from the parent object (GSE227136) using CellTypist ^28^. Distinct populations of immune (myeloid and lymphoid) and non-immune cells (epithelial, endothelial, and fibroblasts) were identified in the whole-lung RNA-seq dataset. Cell-cell communication between lymphocytes (CD4⁺, CD8⁺/NKT-like cells, and NK cells) and epithelial cells [alveolar type 1 (AT1), AT2, transitional AT2, KRT5⁻/KRT17⁺, and secretory cells] was assessed using LIANA and cell-to-cell tensor ^29^. In the CITE-seq dataset, eleven major T-cell clusters were identified. DESeq2 ^30^ pseudobulk analysis was performed to identify differentially expressed genes among T-cell populations in IPF compared to control samples.

### Statistics

GraphPad Prism software v8 (GraphPad, La Jolla, CA, USA) and unpaired T test were used to determine significant differences between two different conditions.

## Results

### Mapping lymphocyte spatial distribution in IPF lungs using CODEX multiplex tissue imaging

To determine the spatial relationship between T-cells and AECs in IPF, we used CODEX multiplexed tissue imaging to examine the distribution of CD4⁺ and CD8⁺ T cell subsets in IPF lungs. Tissue microarray (TMA) sections were generated from two distinct regions of paraffin-embedded lungs from 6 non-diseased controls and 6 IPF patients (**Supplemental Table 1**). A panel of 14 canonical and functional markers (**Supplemental Table 2**) was used to identify immune cell subsets, epithelial cells (pan-cytokeratin⁺), and endothelial cells (CD31⁺). All subsets of T-cells (CD4⁺ T-cells (CD4⁺), regulatory T cells (T_REG_; FOXP3⁺), and CD8⁺ T cells (CD8⁺)) were localized primarily in the alveolar interstitium of both control and IPF lungs; however, we noted that many T-cells were located in proximity to AECs in IPF (**Fig. 1A–B**). Quantification of T-cell subsets revealed a significant increase in CD4, T_REG,_ and CD8 cell numbers in IPF compared to controls (**Fig. 1C**).

**Figure 1:**
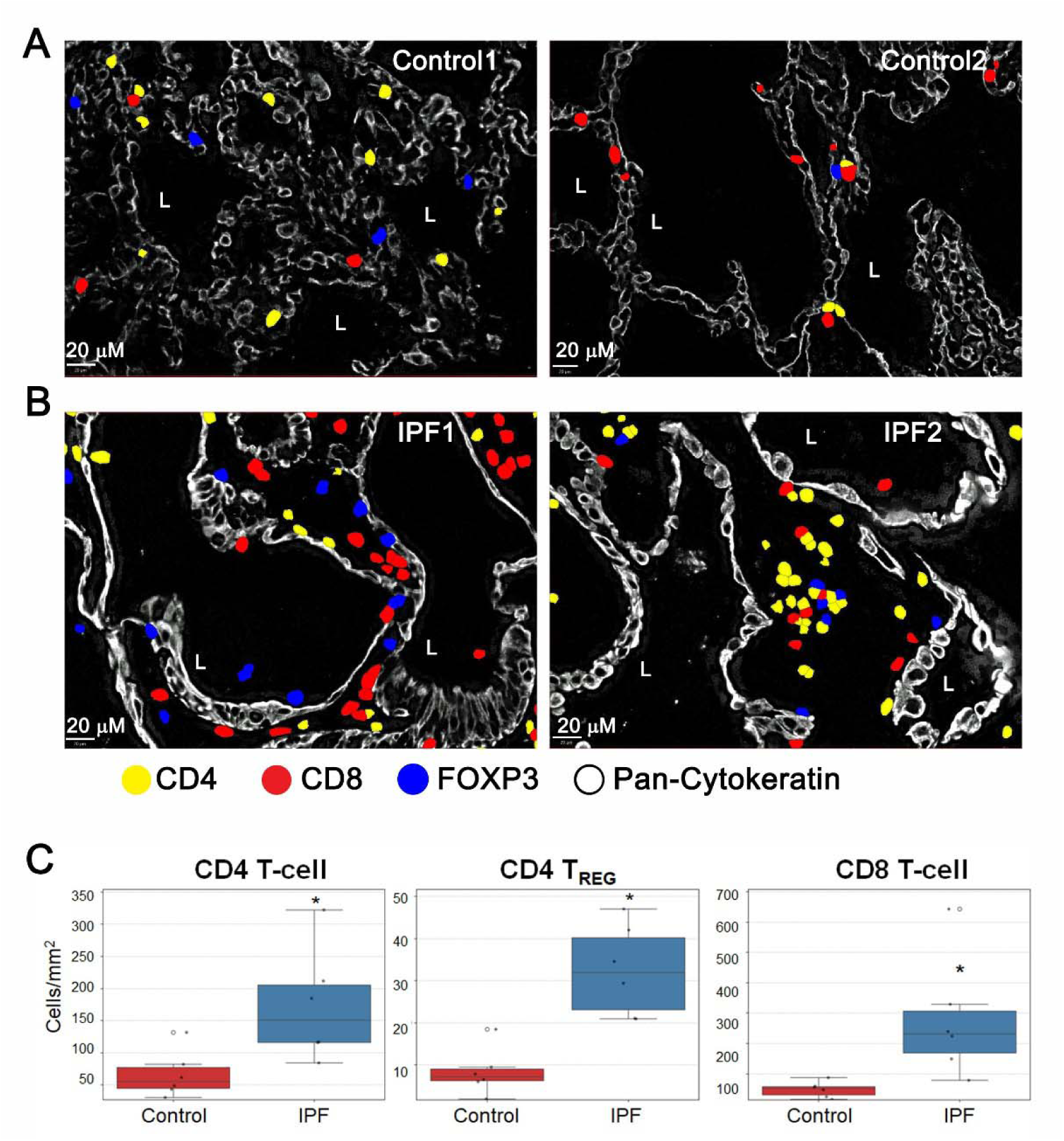
**A–B** Multiplexed spatial imaging using CODEX showing lung sections from two representative control and IPF donors. Images were analyzed using QuPath, with cell types annotated as epithelial cells [pan-cytokeratin^+^ (white)], CD4 T-cells [CD4^+^ (yellow)], CD4 T regulatory cells (Tregs) [FOXP3^+^ (blue)], and CD8 T-cells [CD8^+^ (red)]. **C** Boxplot displaying the median and interquartile range (IQR) of the normalized CD4 T-cell counts per mm² across control donors (n = 6) and IPF patients (n = 6). Individual donor data points are overlaid. Statistical significance was determined using the Mann-Whitney U Test (*p < 0.05)*.

### Characterization of lung T-cell subsets and their activation profile in IPF

Next, we performed Cellular Indexing of Transcriptomes and Epitopes by sequencing (CITE-seq) using the 10x Genomics platform on CD3⁺ T lymphocytes isolated from lung explants of 10 control patients, 9 IPF patients, and 10 non-IPF ILD patients (**Supplemental Table 1**). Minced lung tissues were processed as previously described ^11^, and total CD3⁺ live T-cells were sorted by flow cytometry. Prior to sequencing, cells were incubated with oligo-conjugated antibodies targeting 15 surface markers for CITE-seq as illustrated in figure 2A (**Supplemental Table 3**).

**Figure 2:**
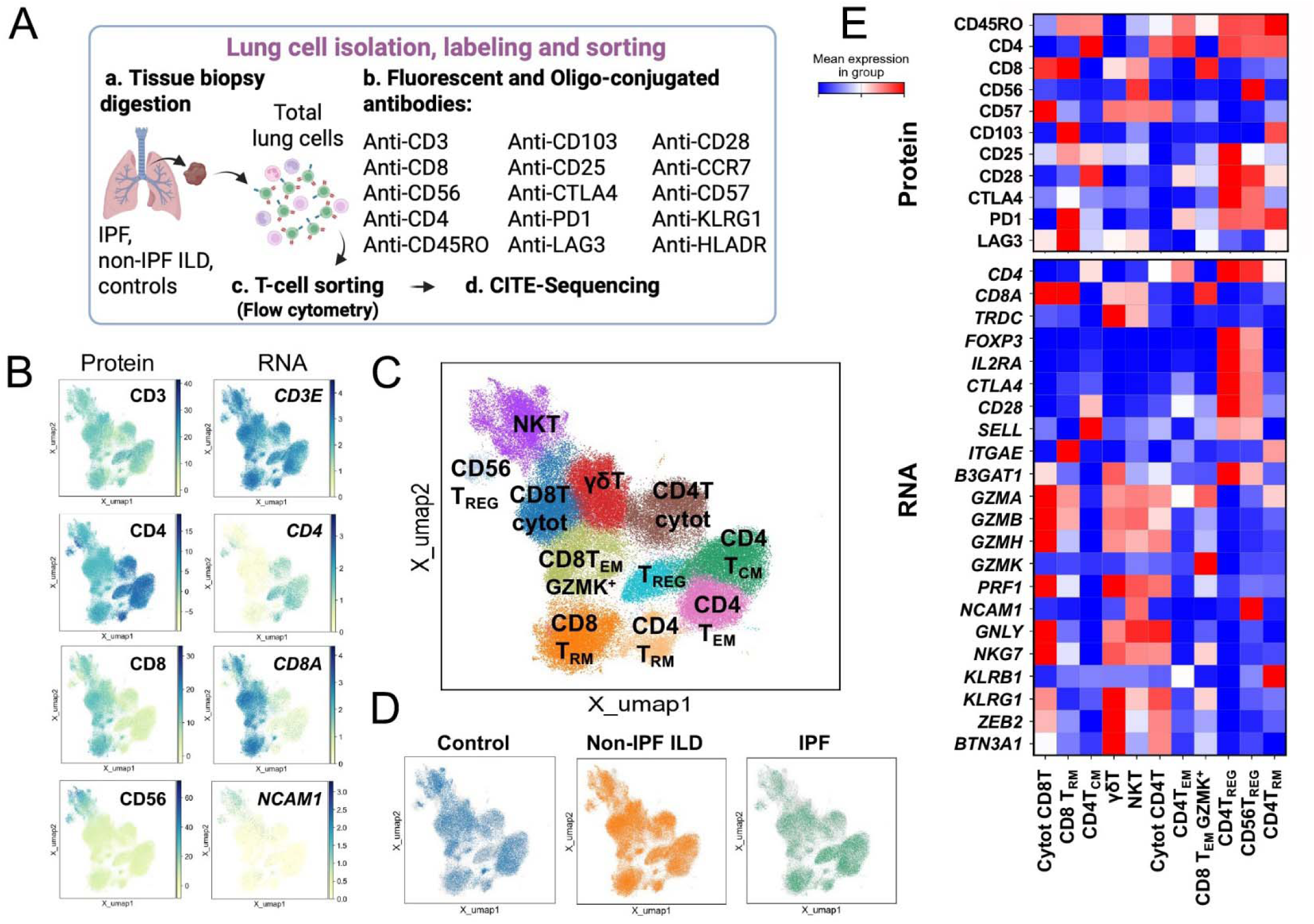
**A** Schematic workflow illustrating the isolation, staining, and sorting of T-cells from lung tissues for CITE-seq analysis. **B** Multimodal cluster identification integrating RNA and protein expression profiles, visualized in Uniform Manifold Approximation and Projection (UMAP) plots from 91,434 cells isolated from ten controls and nineteen fibrotic lungs. **C** UMAP plot showing T-cell subsets annotated based on RNA and protein marker expression, with subsets color-coded as indicated. **D** UMAP plots comparing the distribution of T-cell clusters across controls, non-IPF interstitial lung disease (ILD), and IPF lungs. **E** Heatmap showing protein and RNA expression profiles for signature markers defining each T-cell subset identified using CITE-seq.

Using multimodal analysis of the processed data, we identified eleven subpopulations of CD4⁺ and CD8⁺ T-cells based on combined RNA and protein marker expression (**Fig. 2B**). Clusters were annotated according to canonical T-cell markers as follows: CD4 central memory T-cells (T_CM_; *SELL*⁺), CD4 effector memory T-cells (T_EM_; *CD45RO*⁺*SELL*⁻), CD4 regulatory T-cells (T_REG_; CD25⁺, *FOXP3*⁺), CD4 resident memory T-cells (T_RM_; CD103⁺, *ITGAE*⁺), an unusual population of CD56⁺ T_REG_ cells (CD56⁺, *NCAM1*⁺, *FOXP3*⁺), CD8 T_EM_ (*GZMK*⁺), CD8 T_RM_ (CD103⁺, *ITGAE*⁺), cytotoxic CD4 and cytotoxic CD8 T-cells (*GNLY*⁺, CD56⁻, *NCAM1*⁻), γδ T-cells (*TRDC*⁺), and natural killer-like T-cells (NKT-like; *GNLY*⁺, CD56⁺, *NCAM1*⁺) (**Fig. 2C-E**). Several of these subsets expressed cytotoxicity-related genes, including granzymes (*GZMA, GZMB, GZMH*). All T-cell subsets were identified across participants and within the three groups, though in varied frequencies (**Supplemental Fig. 1A-B**). Next, we utilized our CyTOF dataset from peripheral blood or lungs ^11^ to confirm the existence of CD56⁺ T_REG_ in IPF. CD56 was weakly detected in the lungs, but CD38 (a marker to identify activated NK cells^31^) colocalized with FOXP3 in a subset of immune cells. Thus, we observed a CD56^+^ T_REGs_-like population among CD4^+^CD25^+^ T-cell predominantly in lungs of IPF compared to control donors or peripheral blood (**Supplemental Fig. 1C)**. These findings highlight the presence of a diverse network of T-cell subsets in the lung.

To investigate disease-related changes in gene expression, we performed cell-type specific, subject-level pseudobulk differential expression analyses comparing IPF subjects to controls. Several T-cell subpopulations showed a substantial number of differentially expressed genes (DEGs) in IPF compared to control subsets, with a large majority of DEGs being up-regulated (**Fig. 3A**). CD56 T_REG_ showed the greatest number of DEGs compared to other T-cell subsets. **Figure 3B** shows the top five DEGs for CD4 T_REG_, CD56⁺ T_REG_, CD4 T_EM_, CD4 T_RM_, CD8 T_RM_, and NKT-like cells. In these T-cell subsets, we further examined DEGs using a multivariate linear model and PROGENy [a pathway inference framework based on different pathway resources ^32^] to identify differentially activated pathways. We found that epidermal growth factor receptor (EGFR) signaling, TGFβ, and TNF/Trail signaling were increased in these T-cell subsets, with EGFR signaling consistently being the most up-regulated pathway across subsets (**Fig. 3C**). These results indicate a conserved set of signaling pathways in IPF, which are likely influenced by local factors present in the fibrotic microenvironment.

**Figure 3:**
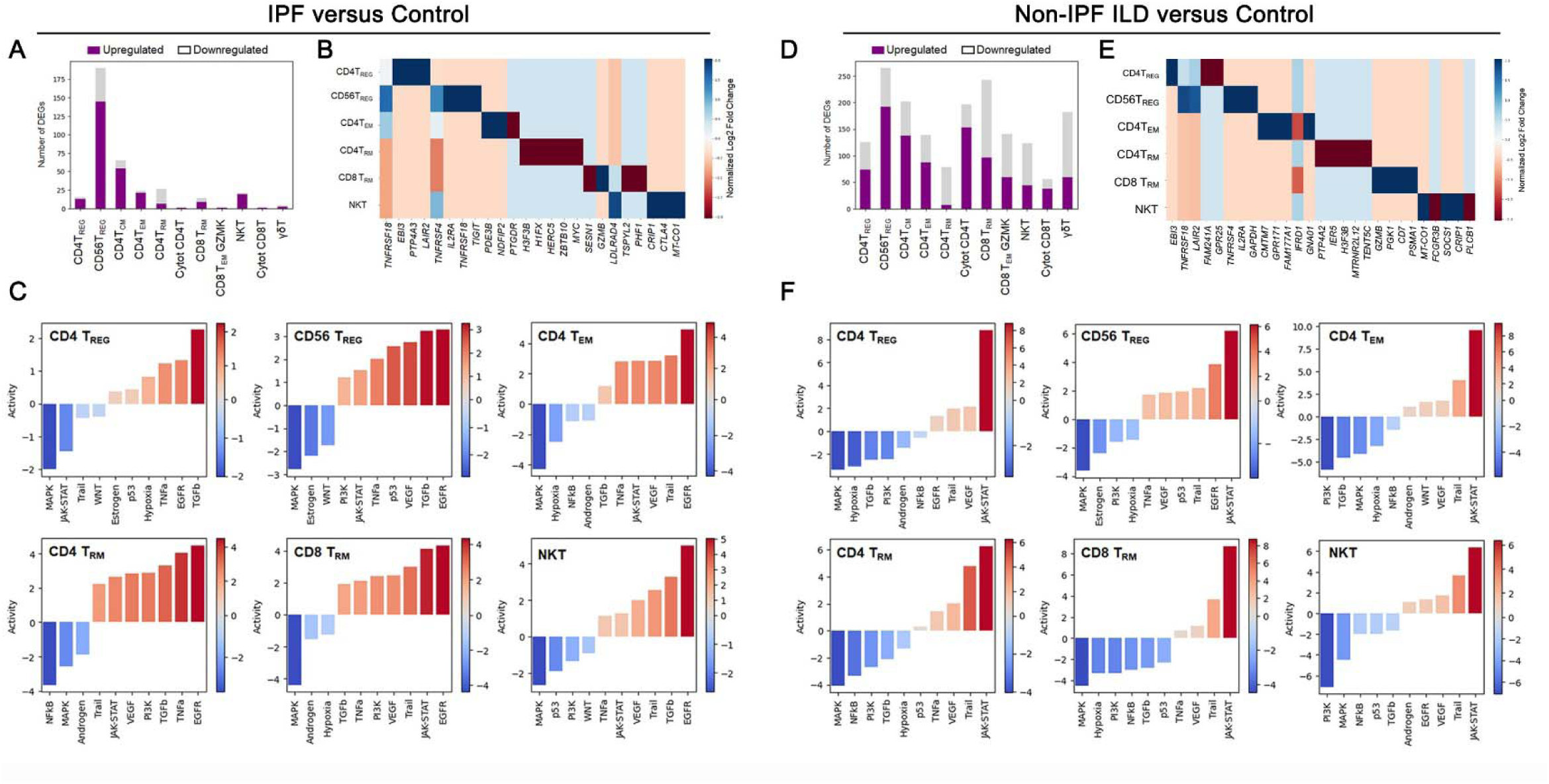
**A.** Bar chart showing the number of upregulated (purple) and downregulated (light gray) differentially expressed genes (DEGs) across T-cell subsets in IPF lungs compared to controls. DEGs were identified using PyDEseq2 pseudobulk analysis, with thresholds of adjusted p-value (padj < 0.05) and log2 fold change > ±1. **B** Heatmap displaying the top five DEGs identified through pseudobulk analysis of T-cell subsets in IPF lungs compared to controls. **C** Barplot showing pathway activity scores for the top ten pathways in six different T-cell subset from IPF lungs, calculated using multilevel modeling (MLM) analysis. MLM was performed on pseudobulk gene expression profiles using the PROGENy database. Pathways were ranked by activity levels, with active pathways shown in red and inactive pathways shown in blue. **D** Bar chart showing the number of upregulated (purple) and downregulated (light gray) DEGs across T-cell subsets in non-IPF ILD lungs compared to controls. DEGs were identified using PyDEseq2 pseudobulk analysis, with thresholds of adjusted p-value (padj < 0.05) and log2 fold change > ±1. **E** Heatmap displaying the top five DEGs identified through pseudobulk analysis of T-cell subsets in non-IPF ILD lungs compared to controls. **F** Barplot showing pathway activity scores for the top ten pathways in six different T-cell subset from non-IPF ILD lungs, calculated using MLM. MLM was performed on pseudobulk gene expression profiles using the PROGENy database. Pathways were ranked by activity levels, with active pathways shown in red and inactive pathways shown in blue.

As a comparison, we analyzed T-cell DEGs in non-IPF ILD subjects compared to control samples. We observed that DEGs were identified in all subsets of T-cells in non-IPF ILD compared to controls, with the majority of DEGs being up-regulated (similar to IPF) (**Fig. 3D**). In addition to finding more DEGs in non-IPF ILD compared to IPF, the top five DEGs in in non-IPD ILD subjects were different from those identified IPF patients (**Fig. 3E**). Importantly, up-regulated pathways in CD4 T_REG_, CD56⁺ T_REG_, CD4 T_EM_, CD4 T_RM_, CD8 T_RM_, and NKT-like cells were different than IPF. Janus Kinase and Signal Transducers and Activators of Transcription [JAK-STAT), TRAIL, and VEGF pathways were upregulated in all these T-cell subsets in non-IPF-ILD compared to controls (**Fig. 3F**). Together, these results indicate important differences in the microenvironment of non-IPF ILD compared to IPF, driving different sets of disease-specific pathways in T-cells.

### Molecular drivers of cell-to-cell communications between T-cells and epithelial cells

To investigate how T-cell/epithelial cell crosstalk may influence pulmonary fibrosis, we analyzed scRNA-seq data from CD45^-^ (non-hematopoietic) and CD45^+^ (hematopoietic) lung cells generated from the same subjects who had undergone T-cell CITE-seq analysis ^23^. The dataset was integrated using Harmony, resulting in thirty-six unsupervised clusters that we annotated as immune and structural cells, including lymphocytes, lymphactic, myeloid, epithelial, basal, fibroblasts, and endothelial cells (**Fig. 4A**). The lymphocyte cluster included CD4⁺, CD8⁺/NKT-like, and NK cells, and the epithelial cluster was comprised of AT1, AT2, transitional AT2, KRT5⁻/KRT17⁺, and three subsets of secretory cells: SCGB1A1⁺MUC5⁺, SCGB1A1⁺SCGB3A2⁺, and SCGB3A2⁺. To identify the network of ligand-receptor pairs and the most common cell interactions between the different populations of T-cell and epithelial cells, we applied LIANA and Tensor cell-to-cell analysis between lymphocytes and epithelial cells (illustrated in **Fig. 4B**). This analysis combines the LIANA analysis into a 4D tensor to capture the coordinated relationship between ligand-receptor interactions and communicating cell type pairs in multiple samples^29^. As a result, we generated six factor-specific networks representing cell-cell interactions among the epithelial and lymphocyte populations. Factors 1, 4, 5, and 6 revealed connections between different subsets of epithelial cells, while Factors 2 and 3 highlighted interactions between lymphocytes and epithelial cells (**Fig. 4C-D**). Focusing on Factor 2, which shows signals from epithelial cells to lymphocytes, we observed that interactions between AT2 cells and T-cell subsets were stronger than those between AT1 cells and T-cells. Additionally, transitional AT2 cells and KRT5⁻/KRT17⁺ cells (abundant cell types found in fibrosis) also showed strong interactions with both CD4⁺ and CD8⁺/NKT-like cells (**Fig. 4E**). In Factor 3 (signals from lymphocytes to epithelial cells), we observed robust communication between CD4 and NK cells with KRT5⁻/KRT17⁺ cells (**Supplemental Fig. 2A**) To determine specific ligand–receptor pairs in these interactions, we explored a bipartite network plot from Factors 2 and 3 and identified the chemokine macrophage migration inhibitory factor (MIF), which is a ligand for CXCR4 and CD74 on T-cells (**Fig. 4F**). These interactions could be important for the recruitment and activation of T-cells as well as macrophages.

**Figure 4:**
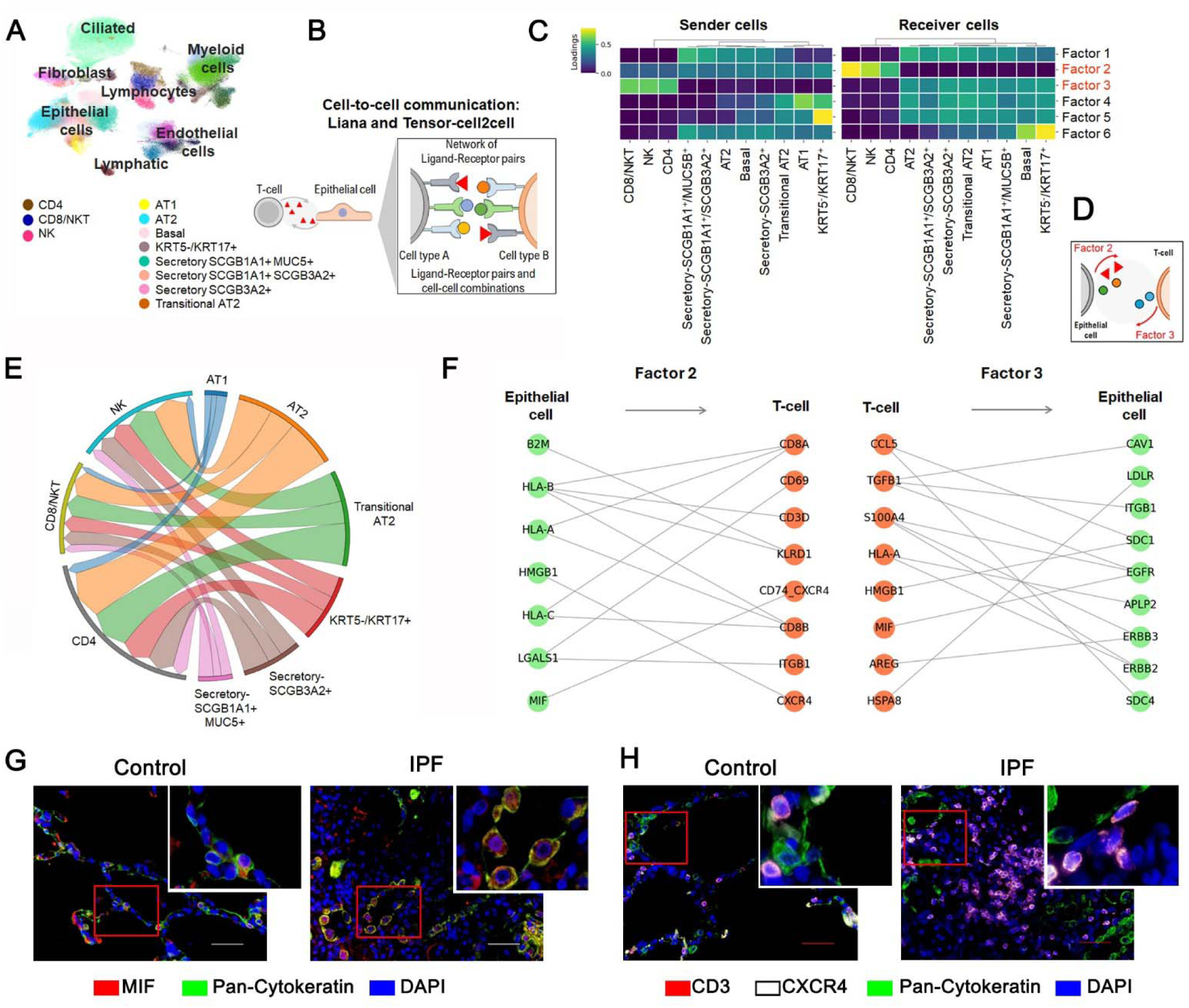
**A** Uniform Manifold Approximation and Projection (UMAP) plot showing whole lung cell populations derived from 224,404 cells isolated from 17 fibrotic lungs and 10 control lungs. **B** Schematic workflow illustrates the computational analysis of ligand-receptor networks and cell-cell interactions using LIANA and Tensor-cell2cell. **C** Heatmap displaying the sender and receiver cell populations during cell-to-cell interactions between the indicated T-cell/NK subsets and epithelial populations using LIANA and Tensor-cell2cell. **D** Illustration of factors 2 and 3, highlighting sender and receiver cell populations during epithelial/T-cell communication. **E** Circos plot visualizing the relationships between epithelial subsets and T-cell subsets based on factor 2. **F** Bipartite network plot visualizing significant ligand-receptor interactions associated with Factors 2 (epithelial to T-cells) and 3 (T-cell to epithelial cell). Nodes represent ligands and receptors, while edges indicate interactions with weights greater than 0.005. **G–H** Immunofluorescence images of lung tissue microarray (TMA) sections from 6 controls and 6 IPF patients: (**G**) staining for macrophage migration inhibitory factor [(MIF), red), pan-cytokeratin (green), and DAPI (dark blue); (**H**) staining for CD3 (red), CXCR4 (white), pan-cytokeratin (green), and DAPI (dark blue).

Transcripts for MIF were expressed especially by transitional AT2 and KRT5^-^KRT17^+^ epithelial cells (**Supplemental Fig. 2B**). Epithelial cells also expressed varied levels of human leukocyte antigen (HLA) class I and II, indicating that these cells could function as antigen-presenting cells to both CD8 and CD4 T-cells (**Supplemental Fig. 2B**). Interactions among different epithelial cells populations are depicted in Supplemental Fig. 2C-F, and showed a variety of ligand/receptor pairs, including adhesion molecules (ICAM1), members of the tissue inhibitor of metalloproteinase (TIMP) family, TGFβ and MMP7, corroborating with an intense tissue remodeling process. We performed an immunofluorescence staining assay for MIF and pan-cytokeratin (**Fig. 4G**) or CD3, CXCR4 and pan-cytokeratin (**Fig. 4H**) in the lung sections of control and IPF lungs. We observed that MIF is found intracellularly in pan-cytokeratin^+^ cells (epithelial cells). Moreover, CXCR4^+^ T-cells were found especially abundant in the interstitium of IPF lungs. Together, these results indicate that epithelial cells may reshape the local immune environment by communicating with T-cells via the MIF-CXCR4-CD74 axis. Because CXCR4/CD74 and EGFR signaling share several downstream intracellular signals and exhibit crosstalk mechanisms ^33, 34^, our data suggest that these receptors might engage in crosstalk during T-cell activation in IPF. We examined the expression of EGFRs, CXCR4 and CD74 in T-cell subsets using our CITE-seq analysis. The average log expression of ERBB2 and ERBB3 was lower compared to CXCR4 and CD74 (**Supplemental Fig. 3**). However, we observed that ERBB2 was detected at high levels in a small group of cells in all populations, but in greater abundance in NKT-like cells. Therefore, the interplay between EGFR, CXCR4, and CD74 signaling pathways may represent a significant cooperative mechanism driving the activation of several population of T-cells in IPF lungs.

### Spatial distribution of functional T-cell subsets indicates a suppressive and cytotoxic environment in IPF lungs

Our findings suggest that CD56 T_REGs_ are a rare and distinctive T-cell population in IPF lungs. To further investigate the presence of additional T-cell subsets in lung tissue, we leveraged our CODEX imaging dataset, which includes markers for CD56, FOXP3, and granzyme B (GZMB). We first annotated CD8 GZMB^+^ T-cells and CD56 T_REGs_ and analyzed the spatial organization of all T-cell populations relative to the nearest epithelial cell (including CD4, CD8, and T_REGs_) using controlsand IPF lung sections. A cumulative distribution plot revealed that CD4 T-cells, CD8 GZMB^+^ T-cells, and T_REGs_ were the closest populations to epithelial cells, whereas other CD8 T-cells and CD56 T_REGs_ were positioned at a greater distance from epithelial cells (**Fig. 5A**). Figure 5B shows a representative image of the spatial location of CD8 GZMB^+^ T-cells in lung tissue from one control and one IPF donor. The number of CD8 GZMB^+^ T-cells was similar between the two groups analyzed (**Fig. 5C**). The spatial distribution of T_REGs_ and CD56 T_REGs_ is shown in Figure 5D, and the counts of CD56 T_REGs_ were significantly increased in IPF lungs compared to control lungs (**Fig. 5E**). We also observed that CD56 T_REGs_ exhibited cytoplasmic localization of the transcription factor FOXP3, whereas conventional T_REGs_ expressed FOXP3 exclusively in the intranuclear compartment (**Fig. 5D**).

**Figure 5:**
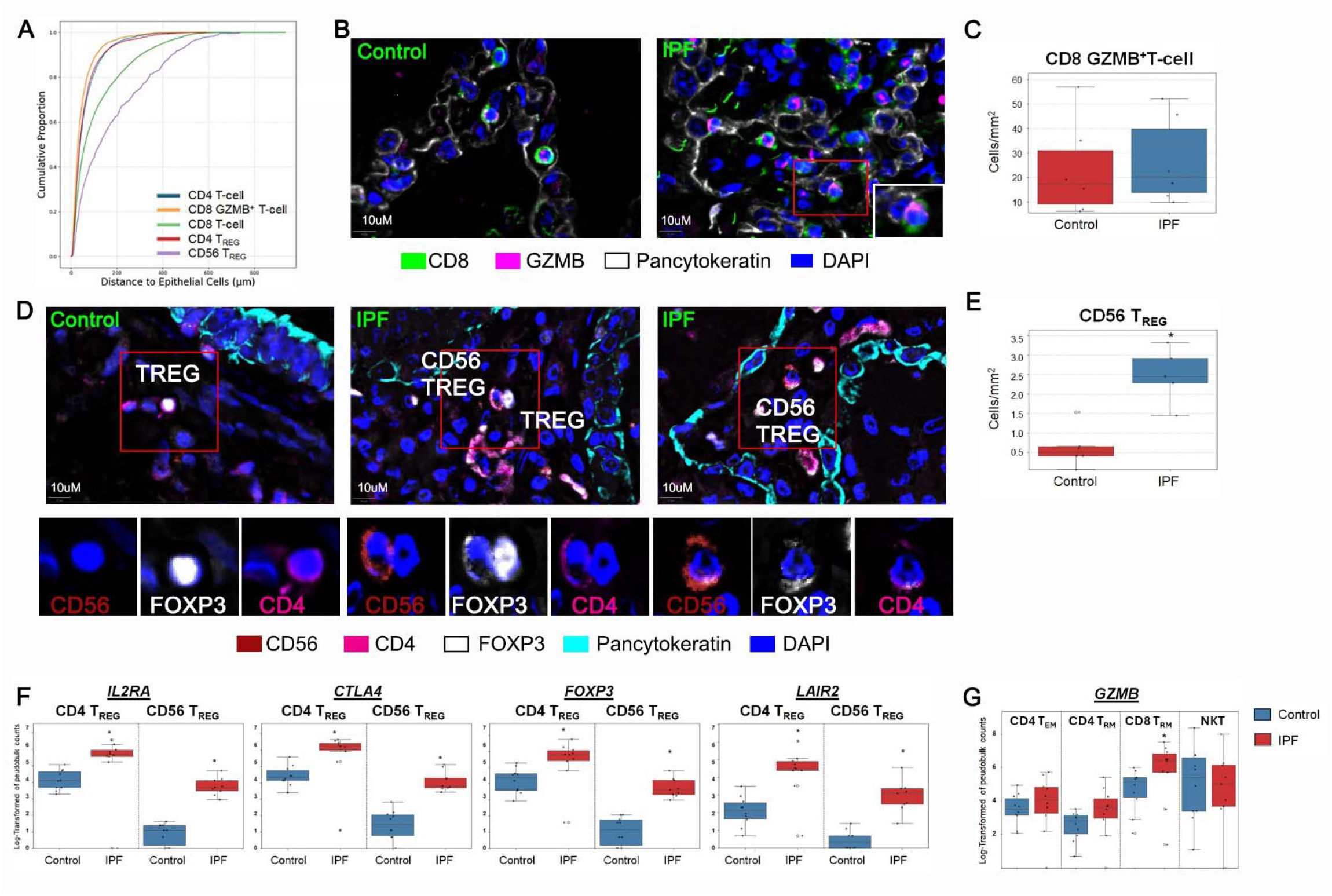
**A** Cumulative distribution plot showing the distances between the indicated T-cell subsets and the nearest epithelial cells in multiplexed spatial imaging of control and IPF lung tissues. Cell types were annotated using QuPath as epithelial cells (pan-cytokeratin^+^), CD4 T-cells (CD4^+^), CD4 T regulatory cells (Tregs, FOXP3^+^), CD8 T-cells (CD8^+^), CD8 GZMB^+^ T-cells (CD8^+^ GZMB^+^), and CD56 Tregs (CD56^+^FOXP3^+^). **B** Representative CODEX image showing CD8 GZMB^+^ T-cells in lung tissue microarray (TMA) sections from control and IPF donors, stained for CD8 (green), granzyme B (GZMB, pink), pan-cytokeratin (white), and DAPI (blue**). C** Boxplot displaying the median and interquartile range (IQR) of normalized CD8 GZMB^+^ T-cell counts per mm² across control donors (n = 6) and IPF patients (n = 6). Individual donor data points are overlaid. **D** Representative CODEX image showing CD4 T_REG_ and CD56 T_REG_ in TMA sections from control and IPF donors (two distinct areas), stained for CD56 (green), FOXP3 (white), CD4 (pink), pan-cytokeratin (aqua), and DAPI (dark blue). **E** Boxplot displaying the median and interquartile range (IQR) of normalized CD56 T_REG_ counts per mm² across control donors (n = 6) and IPF patients (n = 6). Individual donor data points are overlaid. **F** Boxplot displaying the median and interquartile range (IQR) of log-transformed expression levels of IL2RA, CTLA4, FOXP3, and LAIR2 in CD4 T_REG_ and CD56 T_REG_ from control and IPF donors, based on pseudobulk T-cell CITE-sequencing data. **G** Boxplot displaying the median and interquartile range (IQR) of log-transformed GZMB expression levels in CD4 T_EM_, CD4 T_RM_, CD8 T_RM_, and NKT-like cells from control and IPF donors, based on pseudobulk T-cell CITE-sequencing data. Statistical significance was determined using the Mann-Whitney U Test (p < 0.05).

Since FOXP3 controls the expression of various genes involved in immunosuppression, including CTLA4 (an inhibitory molecule) and IL2RA (a high-affinity receptor for IL-2), we compared FOXP3 RNA along with these two markers in T_REGs_ and CD56 T_REGs_ from controls and IPF cells using our CITE-seq dataset. We observed that T_REGs_ exhibited higher levels of IL2RA, CTLA4, and FOXP3 compared to CD56 T_REGs_, but all markers were significantly upregulated in IPF cells relative to controls for both cell types (**Fig. 5F**). Additionally, we observed that both IPF T_REGs_ and CD56 T_REGs_ expressed higher levels of LAIR2 (Leukocyte-Associated Immunoglobulin-Like Receptor 2) compared to controls. LAIR2 is a soluble decoy receptor that binds to collagen and has been shown to be expressed by FOXP3-expressing cells ^35^. Thus, these results indicate that T_REGs_ have conserved functions in IPF, and the immunosuppressive environment could favor the surge of CD56 T_REGs_ with a similar transcriptional program as conventional T_REGs_. Finally, we examined the levels of *GZMB* in memory T-cells, and we observed that CD8 T_RM_ in IPF exhibited higher *GZMB* compared to control cells. Together, these findings show that immunosuppressive and cytotoxic responses are activated by different functional subsets of T-cells present in the lungs of patients with IPF. The proximity and interactions of T-cells with epithelial cells have the potential to influence the survival and differentiation of epithelial cells that favor disease progression.

## Discussion

In this study, we provide a comprehensive characterization of T-cell phenotypes and their spatial distribution within the lungs of IPF patients, shedding light on their potential mechanisms of action influencing pulmonary fibrosis. Our findings reveal that the predominant localization of T-cells within the interstitium may facilitate direct interactions and crosstalk with epithelial cells.

We identified a rare CD56^⁺^ T subset that is highly activated in pulmonary fibrosis and exhibits a strong immunosuppressive phenotype. Furthermore, we highlight the MIF/CXCR4-EGFR signaling axis as a significant pathway mediating epithelial-T-cell interactions in the fibrotic lung environment. Collectively, these results underscore the relationship between T-cell subsets and dysfunctional epithelial cells, such as transitional AT2 cells and KRT5^-^/KRT17^+^ cells, and demonstrate how different T-cell subsets can be activated by environmental signals to carry out their specific functions. Understanding these subtle interactions among cells could be harnessed to develop therapeutic interventions to ameliorate the chronic injury of epithelial cells in patients with pulmonary fibrosis.

CITE-seq offers a unique opportunity to identify the diverse T-cell populations prevailing in the lungs of patients with pulmonary fibrosis. The multimodal analysis of concomitant expression of protein and RNA enables high-resolution characterization of cell-phenotypes^12–14^, including rare populations observed in our dataset. Our pseudobulk transcriptomic analyses ^36–40,30^ rigorously quantify differences in T-cell activation between IPF and non-IPF ILDs. In non-IPF ILDs, T-cells displayed an activated phenotype dominated by JAK-STAT signaling, a finding that is consistent with well-recognized cytokine-driven immunopathogenic mechanisms of autoimmune diseases^41^ ^42^. In IPF, we observed a distinct activation program regulated by EGFR signaling and TGFβ.

EGFR responses in T-cells elicit proliferation and activation of diverse effector functions^36–40^, including enhancement of FOXP3 expression ^43,44^. In our findings, CD56 T_REGs_ were the most activated T-cell subset in IPF lungs and were responsive to both EGFR and TGFβ activities, while T_REGs_ were activated mainly by TGFβ. CD56 T_REG_ have been identified in several TGFβ-enriched environments ^45,46^, including hepatocellular carcinoma ^45^, atherosclerosis ^47^ and other diseases ^46,48^. However, our data suggest that, in addition to TGFβ, the generation or functional responses of CD56 T_REGs_ might require aditional stimulation such as the ones elicited by EGFR ligands. We observed that the numbers of CD56 T_REG_ were increased in the lungs (where EGF signals are more prevalent) but remained low in peripheral blood of patients, shedding light on a possible tissue-driven origin of these cells.

In this study, we identified the chemokine MIF produced by epithelial cells as a potential link to the recruitment of CXCR4⁺/CD74⁺ T-cells (as previously shown ^49,50^). Elevated MIF levels have been consistently reported in the lungs of IPF patients ^51,52^, and in studies using bleomycin-induced pulmonary fibrosis models, MIF inhibition has beneficial effects on both inflammation (including TGFβ and TNF levels) and tissue injury ^53,54^, emphasizing the significant role of MIF in fibrotic progression. Our scRNA-seq data showed that MIF mRNA was expressed in transitional dysregulated subsets of epithelial cells ^55,56^, which showed robust cell-to-cell interactions with CD4 and CD8 T-cells. Thus, our cell-to-cell interaction analysis, which is consistent with reports that EGFR and CXCR4/CD74 have common intracellular signals and synergistic effects^57^ ^58^ ^59^, indicates that these pathways may culminate in the final activation of T-cells in IPF lungs.

### Importantly, epithelial cells appear as central players in this process

The limitations of our study include that lung tissues were obtained from patients with advanced disease, limiting our ability to determine whether these responses are also present in the early stages of pulmonary fibrosis. Our CITE-seq analysis was performed in different batches to minimize the stress on tissue-isolated cells, which can introduce batch effects in single-cell analysis. To address this, we used subject-level pseudobulk analyses and conservative statistical approaches to mitigate these effects on differentially expressed (DE) genes.

Nonetheless, the larger number of DE genes observed in non-IPF ILD samples may in part reflect some batch-related differences. Additionally, our study lacks the statistical power to segregate patients based on treatments, preventing us from determining the impact of antifibrotic or anti-inflammatory therapies on T-cell functions. Exploring differences in T-cell functions between early and late phases of the disease could provide valuable insights into how disease progression influences local T-cell responses.

Despite these limitations, our findings provide important new insights into the pathogenesis of IPF, highlighting both differences and similarities in T-cell functions between IPF and other forms of ILD driven by autoimmune responses. These studies demonstrate that epithelial cells actively influence lung T-cells through conserved communication pathways, and controling MIF/CXCR4 appeared to be a common avenue to block or impair excessive T-cell responses. A deeper understanding of how these pathways are increased and progress in fibrosis could lead to significant therapeutic and translational opportunities aimed at minimizing the risk of epithelial injury in IPF that leads to disease progression.

