## Supplemental Tables and Figures for "T-cell Multiomic Analysis Identifies Subsets and Mechanisms of Interaction with Epithelial Cells in Idiopathic Pulmonary Fibrosis"

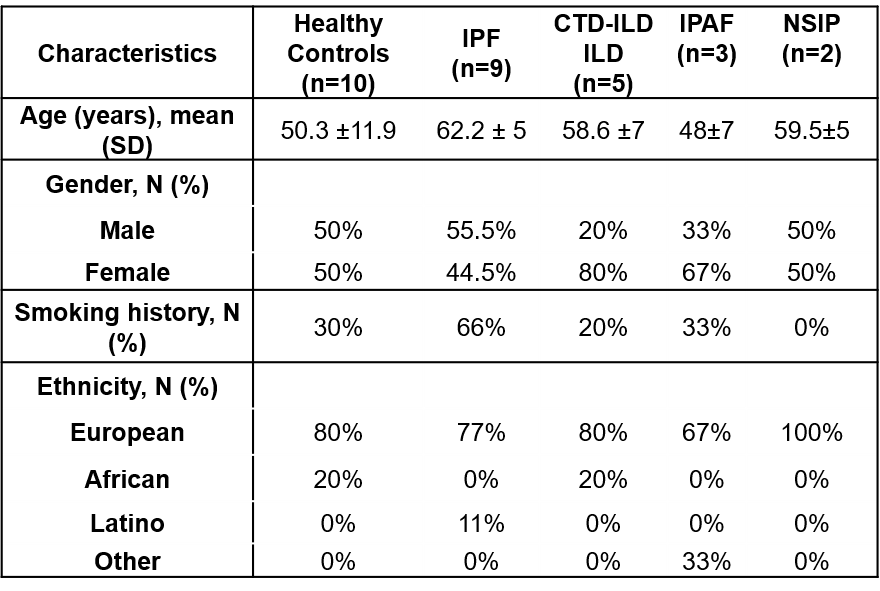

Supplemental Table 1: Characteristics of participants

IPF= Idiopathic Pulmonary Fibrosis

CTD-ILD = connective tissue disease-associated ILD

IPAF= Interstitial pneumonia with autoimmune features

NSIP=nonspecific interstitial pneumonia

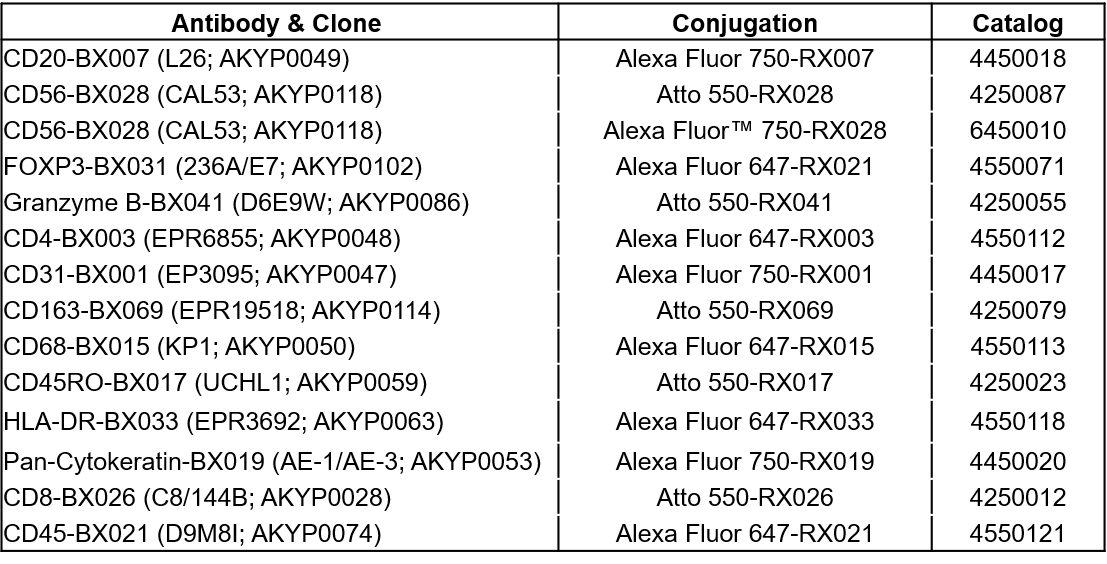

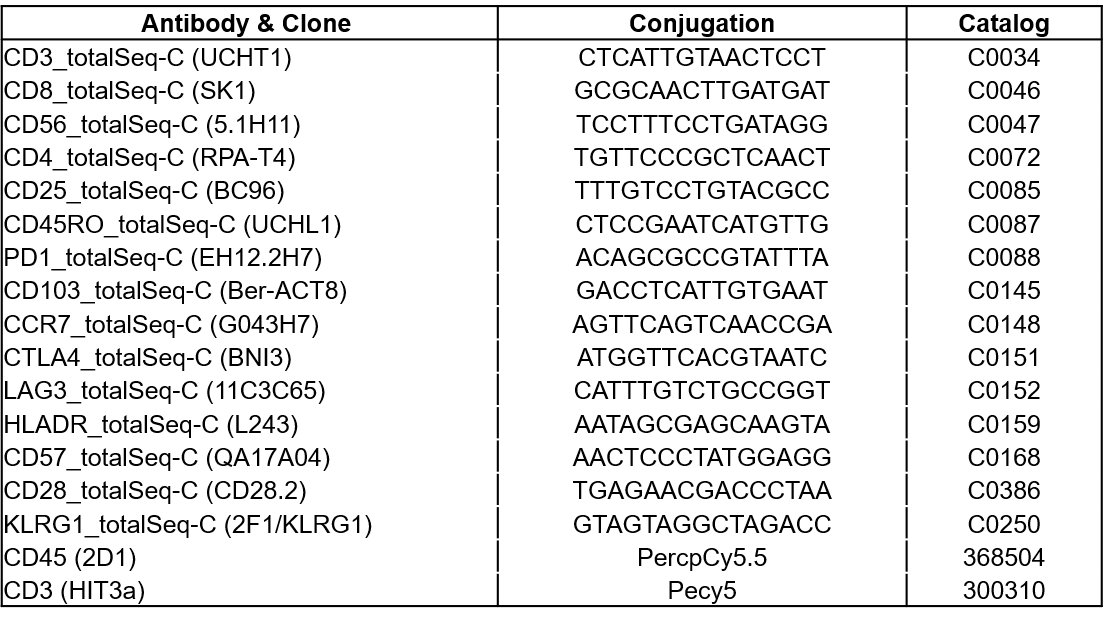

Supplemental Table 2: List of CODEX antibodies

Supplemental Table 3: List of CITE-seq and flow sorting antibody

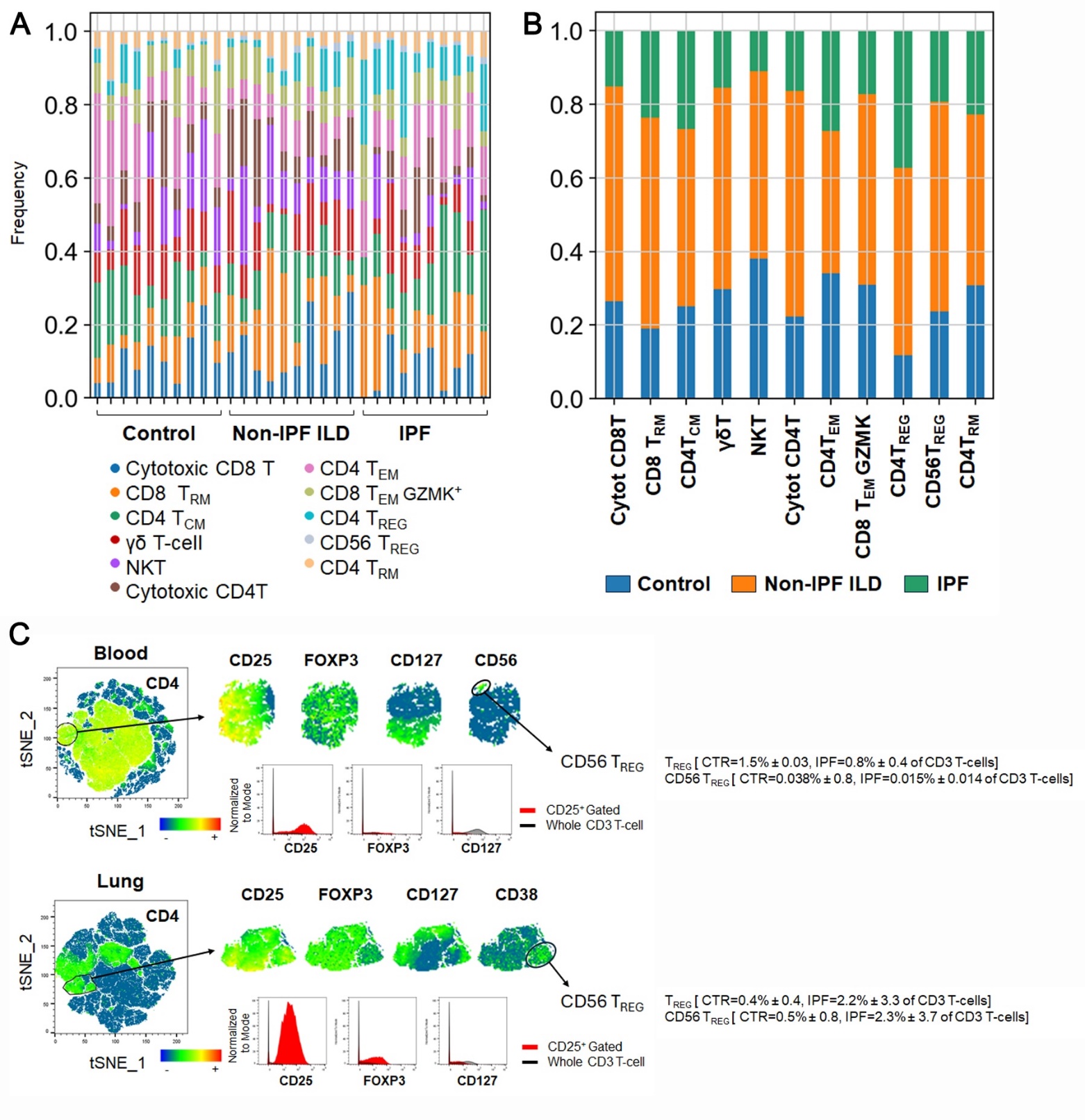

**Supplemental Figure 1:** (**A-B**) Bar charts showing the frequency of the indicated T-cell subsets using CITE-seq analysis in controls, non-IPF ILD, and IPF lungs. (**A**) Data grouped by individual subjects, while (**B**) data grouped by the different conditions. (**C**) viSNE map of CD3^+^ T-cells illustrating the gated CD4^+^CD25^+^ T-cell population in blood and lungs from controls and patients with pulmonary fibrosis. Expression levels of CD25, FOXP3, CD127, CD56, and CD38 in CD4^+^CD25^+^ gated cells are shown. The percentage of CD4^+^CD25^+^ TREGs and CD56^+^ TREGs in controls and IPF individuals is displayed. (n = 45 blood, n = 30 lungs).

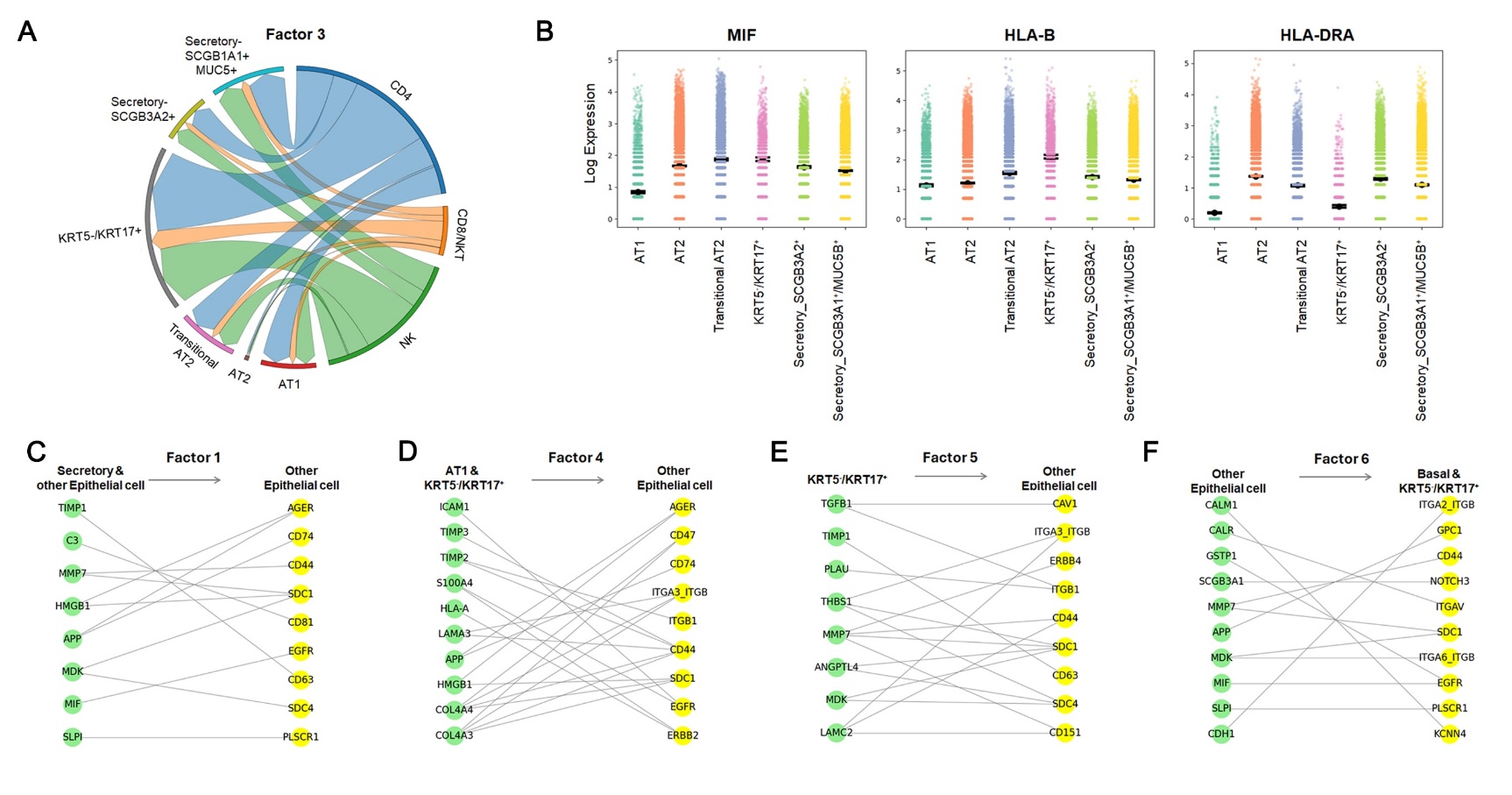

**Supplemental Figure 2: A** Circos plot visualizing the relationships between T-cell subsets and epithelial subsets based on factor 3. **B** Strip and point plot shows log-fold expression of *MIF*, *HLA-B* and *HLA-DRA* in the indicated epithelial cell subsets. **C** Bipartite network plot visualizing significant ligand-receptor interactions associated with Factors 1, 4, 5 and 6 (epithelial to epithelial communication). Nodes represent ligands and receptors, while edges indicate interactions with weights greater than 0.005.

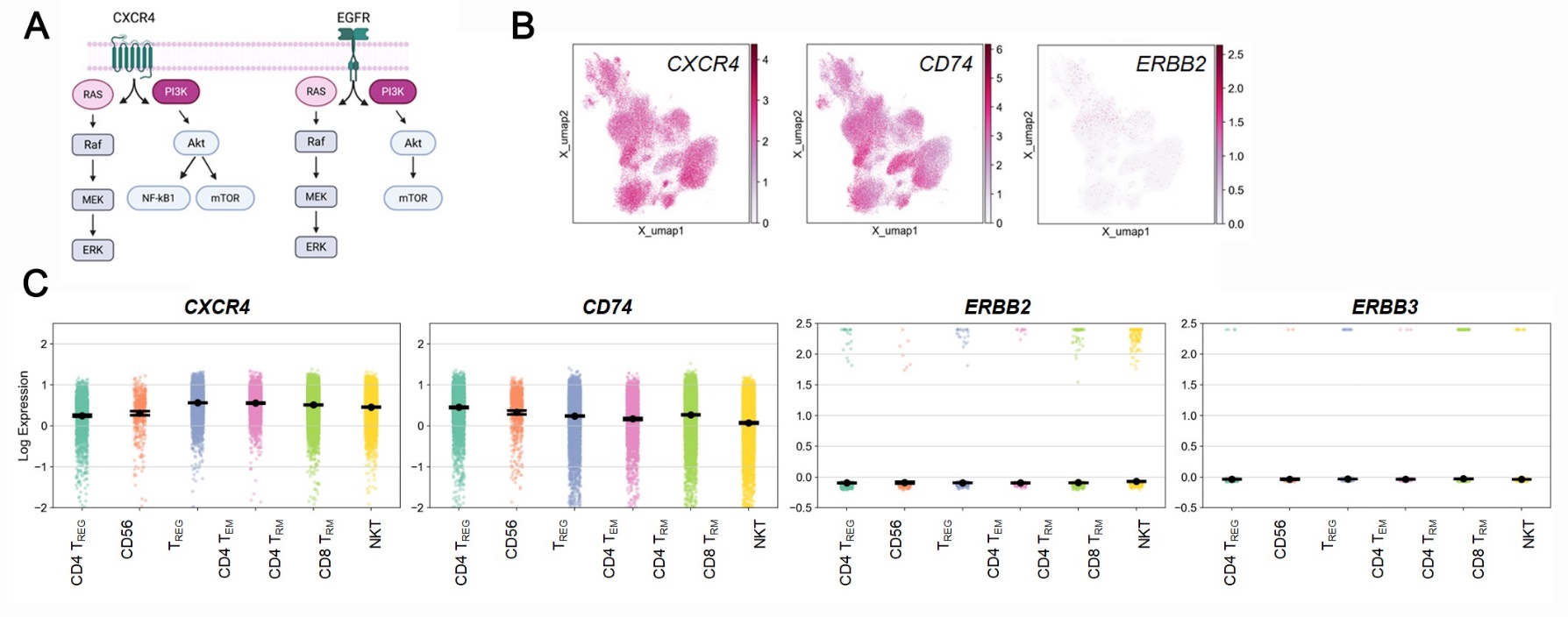

**Supplemental Figure 3: A** Schematic workflow illustrating the intracellular signaling response during CXCR4 and EGFR activation. **B** UMAP plots showing the distribution of CXCR4, CD74, and ERBB across T-cells **C** Strip and point plot shows log-fold expressions of *CXCR4*, *CD74, ERBB2* and *ERBB3* in the indicated T-cell types
